# Determinants of the Divergent *Salmonella* and *Shigella* Epithelial Colonization Strategies Resolved in Human Enteroids and Colonoids

**DOI:** 10.1101/2024.05.03.592388

**Authors:** Petra Geiser, Maria Letizia Di Martino, Ana C. C. Lopes, Alexandra Bergholtz, Magnus Sundbom, Martin Skogar, Wilhelm Graf, Dominic-Luc Webb, Per M. Hellström, Jens Eriksson, Mikael E. Sellin

## Abstract

Despite close relatedness, the major enteropathogens *Salmonella* and *Shigella* differ in infectious dose, pathogenesis, and disease kinetics. The prototype strains *Salmonella enterica* serovar Typhimurium (*Salmonella*) and *Shigella flexneri* (*Shigella*) use Type-3-secretion-systems (T3SSs) to colonize intestinal epithelial cells (IECs), but have evolved partially unique sets of T3SS effectors and accessory virulence factors. A synthesis of how these differences impact the temporal progression of infection in non-transformed human epithelia is missing. Here, we followed *Salmonella* and *Shigella* infections of human enteroids and colonoids by time-lapse imaging to pinpoint virulence factor modules that shape the divergent epithelial colonization strategies. By an apical targeting module that integrates flagella and the SPI-4-encoded adhesin system with T3SS, *Salmonella* accomplishes appreciable numbers of apical invasion events, promptly terminated by IEC death, and thus fostering a polyclonal iterative epithelial colonization strategy. The lack of a corresponding module in *Shigella* makes this pathogen reliant on external factors such as preexisting damage for rare apical access to the intraepithelial environment. However, *Shigella* compensates for this ineptness by an intraepithelial expansion module, where tight coupling of OspC3-dependent temporal delay of cell death and IcsA-mediated lateral spread enables intraepithelial *Shigella* to outrun the IEC death response, fostering an essentially monoclonal colonization strategy.

## INTRODUCTION

*Salmonella* and *Shigella* are major foodborne enteropathogens colonizing the intestinal lumen and invading the epithelium. Despite being closely related and sharing a number of virulence traits, they exhibit a few important differences in their pathogenesis. While non-typhoidal *Salmonella* displays a broad host range and relatively high infectious dose, and primarily infects the distal small intestine and proximal colon in humans or the cecum in mice, *Shigella* is human-restricted with a very low infectious dose of 10 to 100 bacteria in certain patients and selectively colonizes the colonic epithelium (1–3). *Salmonella* and *Shigella*, such as the commonly used prototype strains *Salmonella enterica* serovar Typhimurium and *Shigella flexneri* (here referred to as *Salmonella* and *Shigella*, respectively), compete with the resident gut microbiota and face the soluble mucus layer loaded with antimicrobials as well as the glycocalyx composed of transmembrane mucins and other glycoproteins attached to the apical intestinal epithelial cell (IEC) surface (3, 4).

*Salmonella* employs flagellar motility to reach the epithelial surface (5–7), and flagella have also been implicated in binding to the host cell membrane (8, 9). An extensive repertoire of fimbrial and non-fimbrial adhesins (10–14) can mediate initial *Salmonella* adhesion to host cells, as can the translocon mounted at the tip of the Type-3-secretion-system-1 (T3SS-1) (15, 16). *Shigella*, on the other hand, is non-flagellated, and although a few putative adhesins have been described (17–19), their role during invasion is more controversial, as alternative adhesion mechanisms have also been proposed (20). Both *Salmonella* and *Shigella* employ their syringe-like T3SS to deliver effectors that trigger bacterial uptake by the host cell (3, 21). While it is generally accepted that *Shigella* invades more efficiently form the basolateral side of IECs (22–24), several mechanisms have been proposed for how the epithelial barrier is penetrated from the apical side by either pathogen (20, 25–29). Subsequently, the T3SS is used to shape and maintain the bacteria’s intracellular niche within IECs. While *Salmonella* mainly coopts the endolysosomal pathway to establish a specialized *Salmonella*- containing vacuole via the action of a second T3SS encoded by *Salmonella* pathogenicity island-2 (SPI-2) (30), *Shigella* quickly escapes the endocytic vacuole and adapts a cytosolic lifestyle (3). The T3SS and its effectors are also involved in vacuolar escape (31), mobilization of host actin for cell-to-cell spread via the actin nucleator IcsA (32–34), and have been linked to suppression of host innate immune signaling (35–39).

One key epithelial innate immune mechanism comprises inflammasomes, cytosolic multiprotein complexes assembled upon detection of pathogen-associated molecular patterns (PAMPs) (40). While a range of inflammasomes are activated by different PAMPs, such as flagellin, the T3SS rod and needle proteins and lipopolysaccharide (41–43), they all converge at the cleavage of inflammatory caspases (Caspase-1, 4, 5 or 11), which results in prompt death and extrusion of infected IECs to restrict intraepithelial pathogen loads (44–51), joint with proinflammatory cytokine secretion (47, 49, 51). Enteropathogens, on the other hand, have developed strategies to inhibit epithelial innate immunity, and for *Shigella*, a number of effectors, e.g. OspC3 (35, 36, 52), IpaH7.8 (38) and IpaH9.8 (37), appear capable of suppressing inflammasome signaling cascades at multiple levels.

Although a myriad of host cell interaction mechanisms has been documented for both *Salmonella* and *Shigella*, a synthesis of the decisive virulence factor modules and host cell responses dictating infection outcome remains to be fully established. The mechanisms and molecular targets of isolated *Salmonella* and *Shigella* virulence factors have been studied extensively in transformed or immortalized cell lines and key phenotypes for effector mutants often verified in animal models. However, cell lines incompletely recapitulate primary cell behavior, particularly when it comes to primary IEC-like architecture and functional cell death responses, and they appear hypersusceptible to enterobacterial colonization (29, 53). Furthermore, translating results to/from a highly complex animal model is far from trivial. Due to its human-restricted nature, the pursuit of a reliable animal model for shigellosis has been challenging (54), and a mouse model reproducing some of the typical human pathogenesis has only been developed recently (44).

Enteroids and colonoids offer non-transformed, purely epithelial, physiologically meaningful infection models, which have recently allowed to recapitulate several key aspects of enterobacterial infection mechanisms and host responses at high resolution (23, 24, 45, 46, 53, 55–61). Of note, these intestinal organoid models have thus far primarily been used for static analyses. Hence, we have an incomplete understanding of how bacterial virulence factors and primary IEC responses interact dynamically across infection stages, and how this interplay differs between clinically significant bacteria. Towards this aim, we here followed human enteroid/colonoid infections by time-lapse imaging to pinpoint virulence effector modules that shape the divergent *Salmonella* and *Shigella* intestinal epithelial colonization strategies. Flagellar motility and the SPI-4-encoded adhesin system together with T3SS-1 form an efficient apical targeting module for *Salmonella*, resulting in numerous IEC invasion events typically abrogated by the prompt induction of cell death, thereby generating fast and iterative cycles of IEC invasion and extrusion. *Shigella* lacks a corresponding apical targeting module, but instead employs an efficient intraepithelial expansion module to compensate for exceptionally rare apical IEC binding and invasion. Here, we found that the concerted actions of the inflammasome-targeting T3SS effector OspC3 and the actin nucleator IcsA allow *Shigella* to laterally outrun the epithelial cell death response just in time, thereby fostering an essentially monoclonal epithelial colonization strategy.

## RESULTS

### Unlike wild-type or non-flagellated *Salmonella*, *Shigella* requires preexisting epithelial damage for apical invasion in microinjected human enteroids and colonoids

To follow *Salmonella* and *Shigella* colonization of intestinal epithelia in real-time, we used comparative enteroid and colonoid microinjections combined with time-lapse microscopy. It is well established that intraepithelial *Salmonella* generally occupies a vacuolar niche, the SCV (62), whereas *Shigella* readily escapes the endocytic compartment to colonize the host cell cytosol (3). These differences in intracellular lifestyle led us to use a vacuolar, SPI-2-driven reporter for *Salmonella* (p*ssaG*-GFP, (48, 63)), but a glucose-6-phosphate transporter-based cytosolic reporter (p*uhpT*-GFP, (64)) for *Shigella*. While the p*ssaG*-GFP reporter has previously been applied for *Salmonella* infections in human enteroids (57), we first verified the p*uhpT*-GFP reporter in a standard Caco-2 model for *Shigella* infection (Fig S1A). Enteroid microinjections with motile *Salmonella* wild-type (wt) revealed efficient IEC invasion at multiple sites all around the enteroid (Fig 1A). Flagella-deficient *Salmonella* Δ*fljBfliC* and *Shigella* wt, on the other hand, accumulated atop the epithelium at the enteroid bottom plane within minutes post injection (p.i.). Abundant *Salmonella* Δ*fljBfliC* invasion foci (GFP+) appeared at the enteroid bottom plane at 2-4 h p.i. (Fig 1B), confirming previous results that *Salmonella* flagellar motility may facilitate, but is not strictly required for successful IEC invasion of microinjected enteroids (57). Notably, however, *Shigella* wt remained completely non-invasive even after 16 h p.i. (Fig 1C). Wondering whether epithelial damage could promote *Shigella* invasion, we used the microinjection needle to introduce small scratches (∼1-2 IECs in size) in the epithelium at the enteroid bottom plane prior to luminal microinjection. *Shigella* deposition in close proximity to the basal, outward-facing side of the enteroids was used as an additional control condition. Strikingly, *Shigella* invaded both damaged and basally exposed enteroids with similar efficiencies, showing that epithelial damage enables *Shigella* colonization from the apical face of the enteroid epithelium (Fig 1D-F). *Shigella* infection of colonoids from another donor via these different routes confirmed that the requirement for epithelial damage or basolateral access for *Shigella* invasion of IECs is not segment- or donor-specific (Fig 1G, Fig S1B-C).

**Figure 1.**
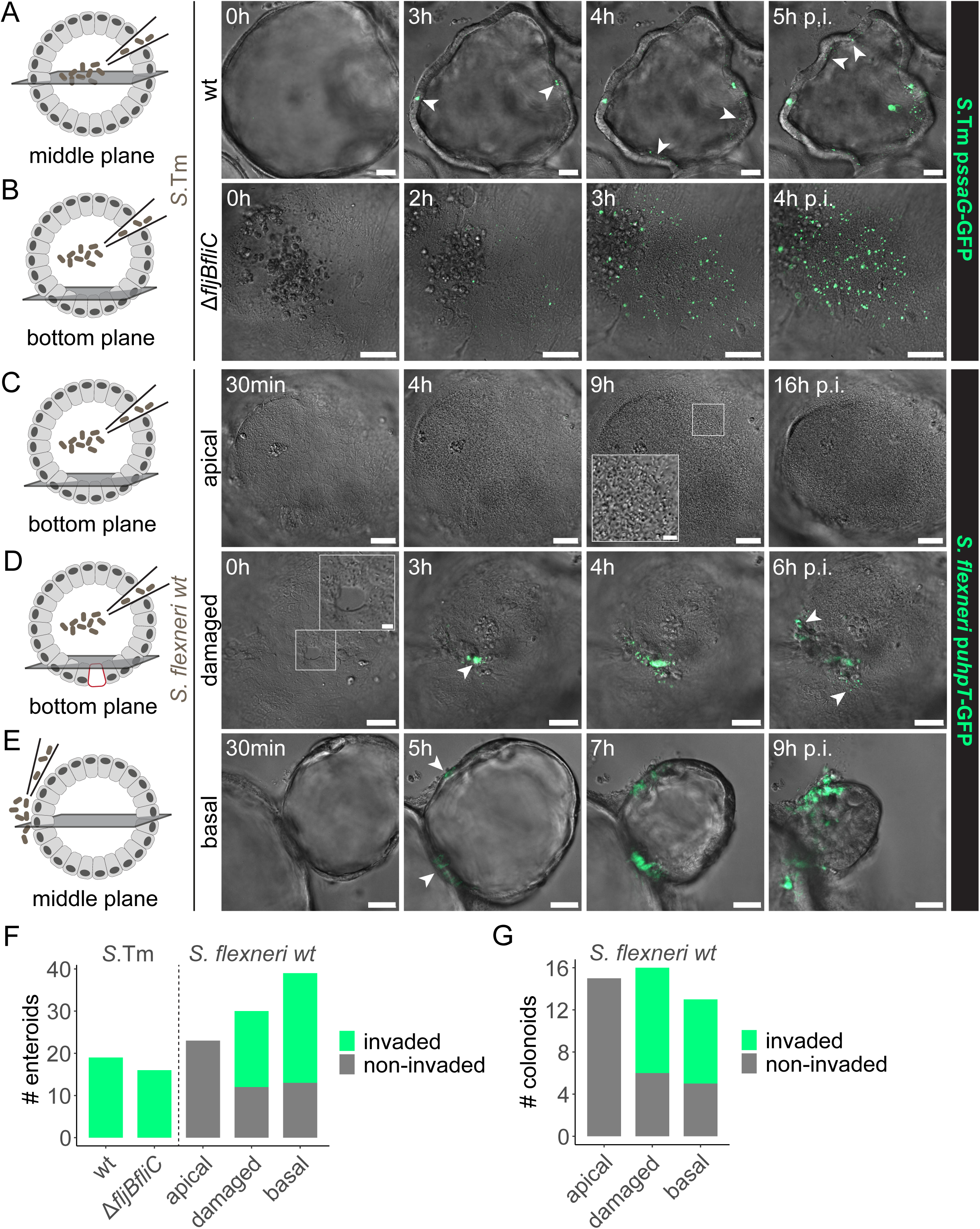
*Salmonella* efficiently colonizes the enteroid and colonoid epithelium from the apical side, but *Shigella* invasion requires epithelial damage. (A-B) Human enteroids were microinjected with (A) *Salmonella* wt and (B) Δ*fljBfliC* harboring the p*ssaG*-GFP intracellular reporter and imaged by wide-field differential interference contrast (DIC) and fluorescence time-lapse microscopy. (C-E) Human enteroids were infected with *Shigella* p*uhpT*-GFP via different routes and imaged as in A-B. (C) Upon luminal microinjection, *Shigella* quickly accumulated at the bottom plane (see insert), but the unperturbed epithelium was not permissive for invasion from the apical side. (D) Epithelial damage introduced with the microinjection needle (see insert) resulted in localized invasion from the site of damage. (E) Basolateral deposition of *Shigella* also resulted in successful epithelial colonization. (F-G) Quantification of the number of (F) enteroids and (G) colonoids invaded by either *Salmonella* strain and *Shigella* wt via the indicated routes of infection. Pooled counts from at least 2 independent experiments are shown. Arrowheads indicate invasion foci. Scale bars: 50 µm (10 µm for inserts). *S*.Tm, *Salmonella* Typhimurium.

### Non-flagellated, SiiE adhesin-deficient *Salmonella* phenocopies *Shigella* in its inability to stably adhere to the apical face of the enteroid and colonoid epithelium

Due to the striking difference in *Salmonella* Δ*fljBfliC* and *Shigella* invasion efficiency at the unperturbed apical face of the epithelium, we next sought to pinpoint the underlying molecular determinants. As a first step towards successful IEC invasion, pathogens have to achieve stable cell surface adhesion. In a newly designed adhesion assay, we tracked Brownian-like motion of constitutively fluorescent (*rpsM*-mCherry, (65)), non-motile bacteria atop the epithelial surface at the bottom plane of microinjected enteroids/colonoids at different time points p.i., anticipating that bacterial Brownian motion would cease completely upon stable surface binding. While non-flagellated *Salmonella* Δ*fljBfliC* displayed two distinct adhering and non-adhering populations already 10-20 min p.i., *Shigella* wt movement did not change over time, and no stably adhering population was observed even at 60-180 min p.i. (Fig 2A-B). Due to the significant difference in invasion event frequency between *Salmonella* and *Shigella* (Fig 1), we also compared adhesion of non-invasive *Salmonella* Δ*fljBfliC* Δ*invG* and *Shigella* Δ*mxiD,* both lacking a structural T3SS (Table S1), finding that the non-invasive strains phenocopied their invasion-competent counterparts (Fig 2C). This confirms that the observed apical adhesion phenotype seen for *Salmonella*, but not to *Shigella*, was not invasion- or T3SS-dependent. Next, we asked what *Salmonella* adhesin(s) might be required to accomplish stable apical IEC binding. Within this string of experiments, enteroids were microinjected with non-motile *Salmonella* Δ*motA ΔSPI-4* (maintains structural flagella (66), but lacks SiiE, a key *Salmonella* adhesin for invasion of polarized epithelia (14)), or non-flagellated *Salmonella* Δ*fljBfliC ΔSPI-4*, in parallel with *Shigella* wt. The absence of SiiE completely abolished stable apical adhesion of either *Salmonella* strain (Fig 2C). Similar results were observed in microinjected colonoids from an independent donor, again with *Salmonella* Δ*fljBfliC ΔSPI-4* phenocopying *Shigella* wt (Fig S2A).

**Figure 2.**
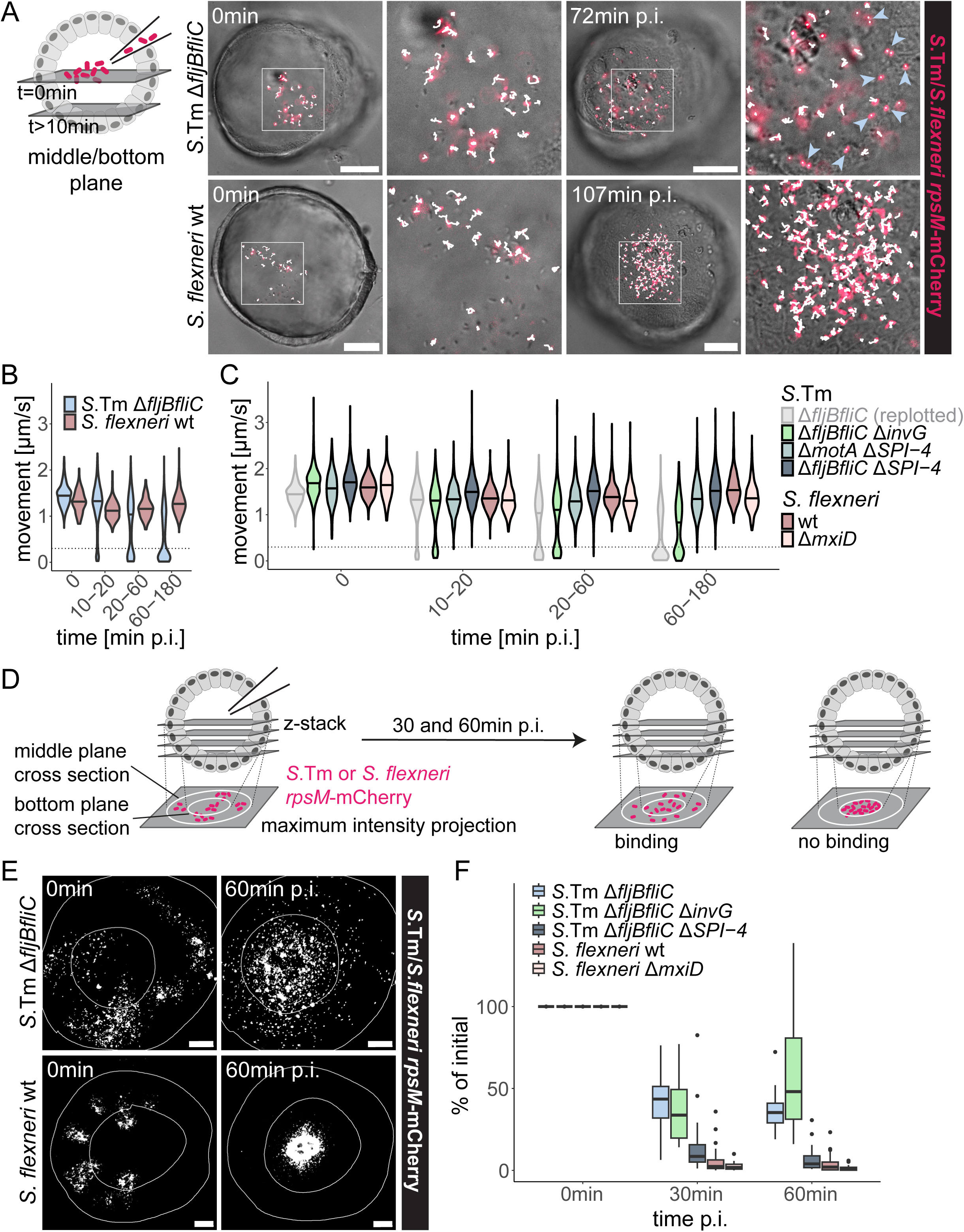
Non-flagellated SPI-4-deficient *Salmonella* and *Shigella* wt fail to stably attach to the apical face of the enteroid epithelium. (A-C) Enteroids were microinjected with the indicated *Salmonella* and *Shigella* strains harboring the constitutive *rpsM*-mCherry reporter, and bacterial movement within the enteroids was tracked at the indicated time points p.i. (A) Schematic of the experimental setup and representative images of bacterial tracking (tracks in white). Adhering bacteria (indicated by light blue arrowheads) were observed for *Salmonella* Δ*fljBfliC* but not *Shigella* wt. Scale bars: 50µm. (B) Quantification of *Salmonella* Δ*fljBfliC* and *Shigella* wt movement speeds at different time points. The bimodal population distribution for *Salmonella* Δ*fljBfliC* with movement speeds close to 0 indicates a stably adhering population. Data from at least 9 enteroids and at least 120 tracks per time point and strain is shown. Horizontal lines indicate the median. (C) Quantification of movement speeds for the indicated *Salmonella* and *Shigella* strains. *Salmonella* Δ*fljBfliC* is replotted from B. Data from at least 8 enteroids and at least 230 tracks per time point and strain is shown. Horizontal lines indicate the median. (D) Schematic of the experimental setup: z-stacks of microinjected enteroids from the middle to the bottom plane were acquired at the indicated time points p.i. Middle and bottom plane cross sections were outlined based on the DIC channel, and bacterial fluorescence (*rpsM*-mCherry) retained at the side epithelium (outer ring), indicative of adhesion, was quantified as mCherry-positive area in maximum intensity projections. (E) In maximum intensity projections with middle and bottom plane outlines, *Salmonella* Δ*fljBfliC* but not *Shigella* wt fluorescence is retained at the side epithelium. Scale bars: 50µm. (F) Quantification of fluorescence retained at the side epithelium relative to the initial mCherry-positive area at the sides. Data from at least 11 enteroids per strain is shown. In the box plots, the height of the boxes represents the interquartile range (IQR), whereas the horizontal line depicts the median. Whiskers extend to the most extreme data point but no further than 1.5x the IQR from the lower (first quartile) or upper (third quartile) boundary of the box. Outliers are indicated as dots. *S*.Tm, *Salmonella* Typhimurium.

In an alternative approach to assess adhesion of non-motile bacteria to the apical IEC surface, we quantified bacterial binding to the internal sides of non-centrally microinjected enteroids. Z-stacks of microinjected enteroids were acquired at different time points p.i. and bacterial fluorescence retained at the enteroid sides was quantified (Fig 2D), hypothesizing that only stable adhesion-competent bacteria would remain at this location, whereas non-binding bacteria should accumulate at the bottom plane. While ∼40% of the initial *Salmonella* Δ*fljBfliC* or *Salmonella* Δ*fljBfliC* Δ*invG* fluorescence was retained at the enteroid sides, *Salmonella* Δ*fljBfliC* Δ*SPI-4, Shigella* wt and *Shigella* Δ*mxiD* accumulated virtually exclusively at the bottom plane with ≤5% of the fluorescence retained at the sides (Fig 2E-F). This finding corroborates our previous results (Fig 2A-C) and emphasizes the SPI-4-encoded SiiE adhesin as the molecular determinant distinguishing non-flagellated *Salmonella* and *Shigella* for stable bacterialadhesion to the apical face of the enteroid epithelium.

In the natural context of motile *Salmonella*, we wondered whether flagellar motility could generate momentum to gain access to the IEC membrane for T3SS-1-dependent stable docking also in the absence of SiiE. To that end, parallel enteroid microinjections with motile *Salmonella* wt and *Salmonella ΔSPI-4* were performed and bacterial movements tracked over time, resulting in three distinct bacterial populations (motile: >5 µm/s, non-motile but exhibiting Brownian motion: 0.3-5 µm/s, and stably bound/invaded: <0.3 µm/s). Notably, *Salmonella ΔSPI-4* still displayed some extent of stable binding/invasion (Fig S2B; 7% vs. 46% *Salmonella* wt at 60-180 min p.i.), while adhesion was abolished in the non-motile strain counterpart (Fig 2C; 0.11% <0.3 µm/s for *Salmonella* Δ*motA* Δ*SPI-4* at 60-180 min p.i.). Hence, the SiiE adhesin (strongly) and flagellar motility (weakly) contribute to make *Salmonella* a highly efficient apical IEC surface binder, in contrast to *Shigella*.

Finally, we quantified bacterial invasion frequencies in 2D enteroid-/colonoid-derived IEC monolayers that allow for elimination of non-invaded bacteria by gentamicin and provide a stable imaging plane over the entire epithelial surface. In line with the results from both adhesion assays, adhesin-proficient *Salmonella* wt and *Salmonella* Δ*fljBfliC* invaded monolayers with relatively high efficiency, while invasion by *Salmonella ΔSPI-4* was reduced (Fig S3A-B). Increased MOI (200) and inclusion of a centrifugation step to force contact with the epithelial surface allowed for rare and unevenly distributed IEC invasion foci also by *Salmonella* Δ*fljBfliC ΔSPI-4* and *Shigella* wt (Fig S3C). Altogether, these results indicate that *Salmonella* harbors an efficient “apical targeting module” hinging on the SPI-4-encoded SiiE adhesin and flagellar motility combined with T3SS-1, which enables IEC entry from the lumen at appreciable frequency. The absence of a corresponding module in *Shigella* makes this bacterium reliant on cooperation between T3SS and external events (e.g. preexisting epithelial damage) to establish an initial foothold in the epithelial lining.

### Efficient expansion of the intraepithelial *Shigella* population compensates for low adhesion and invasion rates, eventually catching up with *Salmonella* epithelial colonization levels

We noted that the vastly higher number of IEC adhesion and invasion events of apical targeting module-proficient *Salmonella* strains was associated with the induction of higher levels of IEC death, as revealed by holes in infected monolayers (Fig S3C). Therefore, the fate of intraepithelial *Salmonella* and *Shigella* populations after successful IEC invasion was further investigated. To that end, we performed time-lapse imaging of individual invasion foci in infected enteroid/colonoid-derived IEC monolayers and included Draq7 as a marker for IEC death and permeabilization (67). As revealed by the abundant Draq7 signal, *Salmonella* wt infection indeed caused prompt induction of IEC death (Fig 3A), often starting even before maturation of the intracellular p*ssaG*-GFP bacterial reporter. By combining a high MOI and centrifugation, occasional IEC invasion events could also be identified and traced for *Shigella* wt and the corresponding “apical targeting module”-deficient *Salmonella* Δ*fljBfliC ΔSPI-4* strain. Notably, while these two strains again phenocopied each other during the early infection stages, *Shigella* wt expanded over time (12-24h p.i.) with dramatically higher efficiency within both enteroid and colonoid, epithelia (Fig 3A-C). Of note, *Shigella* wt induced only minimal levels of IEC death during this intraepithelial expansion phase (Fig 3A). Hence, although apical invasion events are exceptionally rare, *Shigella* (but not *Salmonella* Δ*fljBfliC ΔSPI-4*) compensates for this low adhesion and invasion capacity by efficient post-invasion expansion within the epithelium whilst minimizing the induction of IEC death.

**Figure 3.**
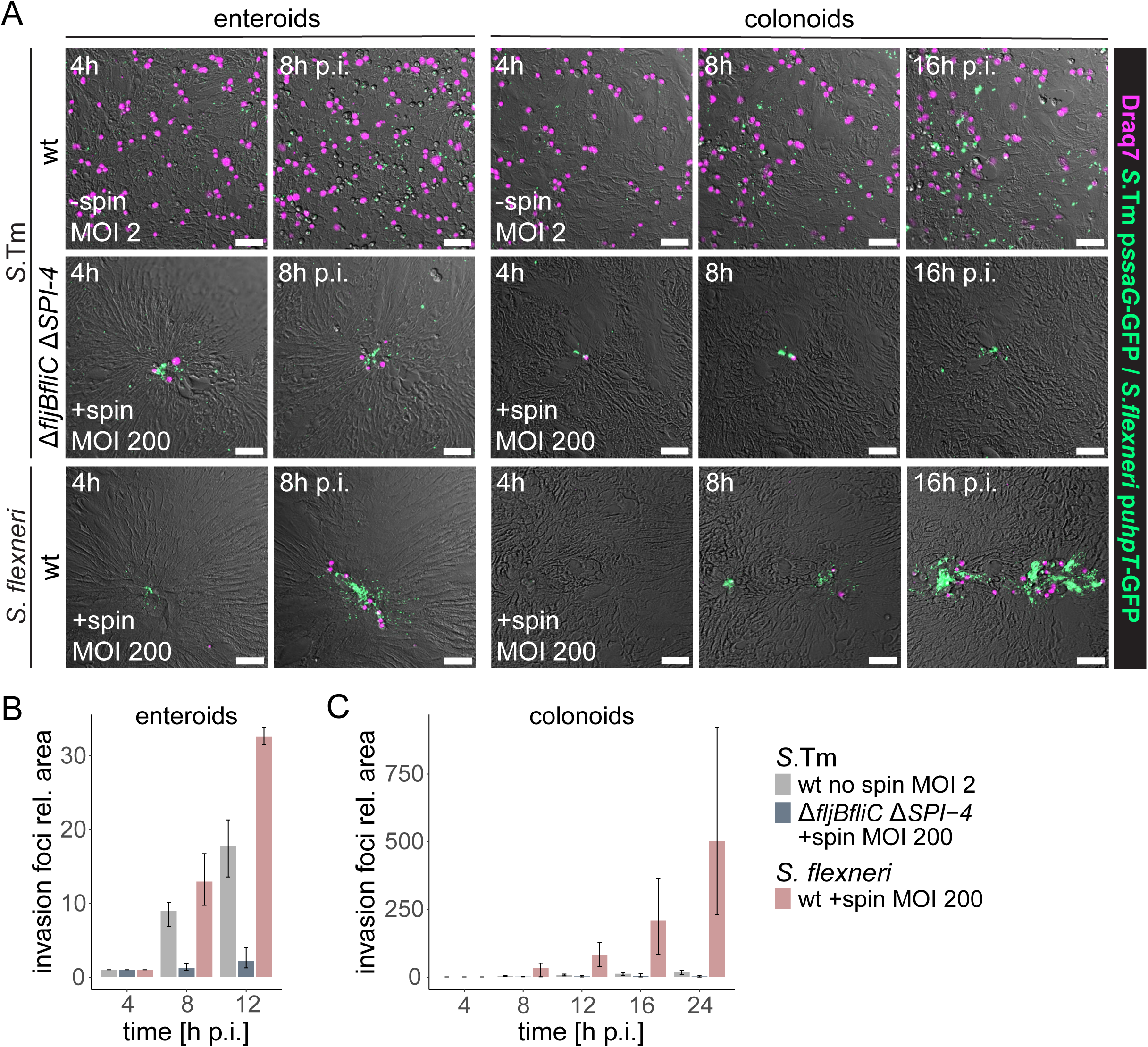
Efficient expansion of the intraepithelial *Shigella* population compensates for rare invasion events. (A-C) Enteroid- and colonoid-derived monolayers were stained with the membrane-impermeable nuclear dye Draq7 and infected with *Salmonella* wt (MOI 2), *Salmonella* Δ*fljBfliC* Δ*SPI-4* and *Shigella* wt (MOI 200 + centrifugation), harboring the intracellular p*ssaG*-GFP or p*uhpT*-GFP reporters, respectively. Individual invasion foci were followed by time-lapse microscopy and their expansion was quantified. (A) Representative images of abundant invasion foci and Draq7-positive cells observed for *Salmonella* wt, rare and confined invasion foci by Δ*fljBfliC* Δ*SPI-4,* and intraepithelial expansion of rare foci with limited Draq7 signal for *Shigella* wt. Scale bars: 50µm. (B-C) Quantification of the GFP-positive area relative to the 4h p.i. time point reveals efficient intraepithelial expansion for *Shigella* in both (B) enteroid- and (C) colonoid-derived monolayers. Data is plotted as mean + range of 3 replicates per strain, with one replicate corresponding to the mean of 6-10 fields of view for an individually infected well.

### The concerted action of OspC3 and IcsA enable *Shigella* to outrun the epithelial inflammasome cell death response by lateral spread

Based on the radically differing fates of intraepithelial *Salmonella* and *Shigella* populations, we next sought to address the interplay between intraepithelial expansion and induction of IEC death, and to pinpoint the *Shigella* virulence factors involved. Within this effort, 2D enteroid/colonoid-derived monolayers were infected with *Shigella* wt, *Shigella* Δ*ospC3* lacking a T3SS effector previously reported to suppress non-canonical (Caspase-4/11) inflammasome activation (OspC3; 35, 36, 52), or *Shigella* Δ*icsA* deficient for the nucleator for cytosolic actin polymerization and cell-to-cell spread (IcsA; 32–34). Infected monolayers were incubated in the presence of Draq7 and individual invasion foci were followed by time-lapse imaging to quantify their expansion and induction of IEC death. While *Shigella* wt spread efficiently within the enteroid epithelium, *Shigella* Δ*ospC3* and *Shigella* Δ*icsA* did not, and such invasion foci were frequently cleared from the epithelium by the prompt induction of cell death (Fig 4A-B). Interestingly, the total Draq7-positive area was broadly comparable for *Shigella* wt and *Shigella* Δ*ospC3* infections (Fig 4C), but since invasion foci differed significantly in size (Fig 4B), the Draq7-to-GFP ratio was analyzed as a measure for the probability of cell death induction per infected IEC. This ratio was markedly higher for *Shigella* Δ*ospC3* than for *Shigella* wt over the entire period of imaging (Fig 4D; 4-20h p.i.). Essentially identical results were obtained also in colonoid-derived monolayers from an independent donor (Fig 4E-G). This shows that OspC3-dependent delay/suppression of IEC death allows sufficient time for IcsA-dependent lateral *Shigella* spread and thereby successful intraepithelial expansion.

**Figure 4.**
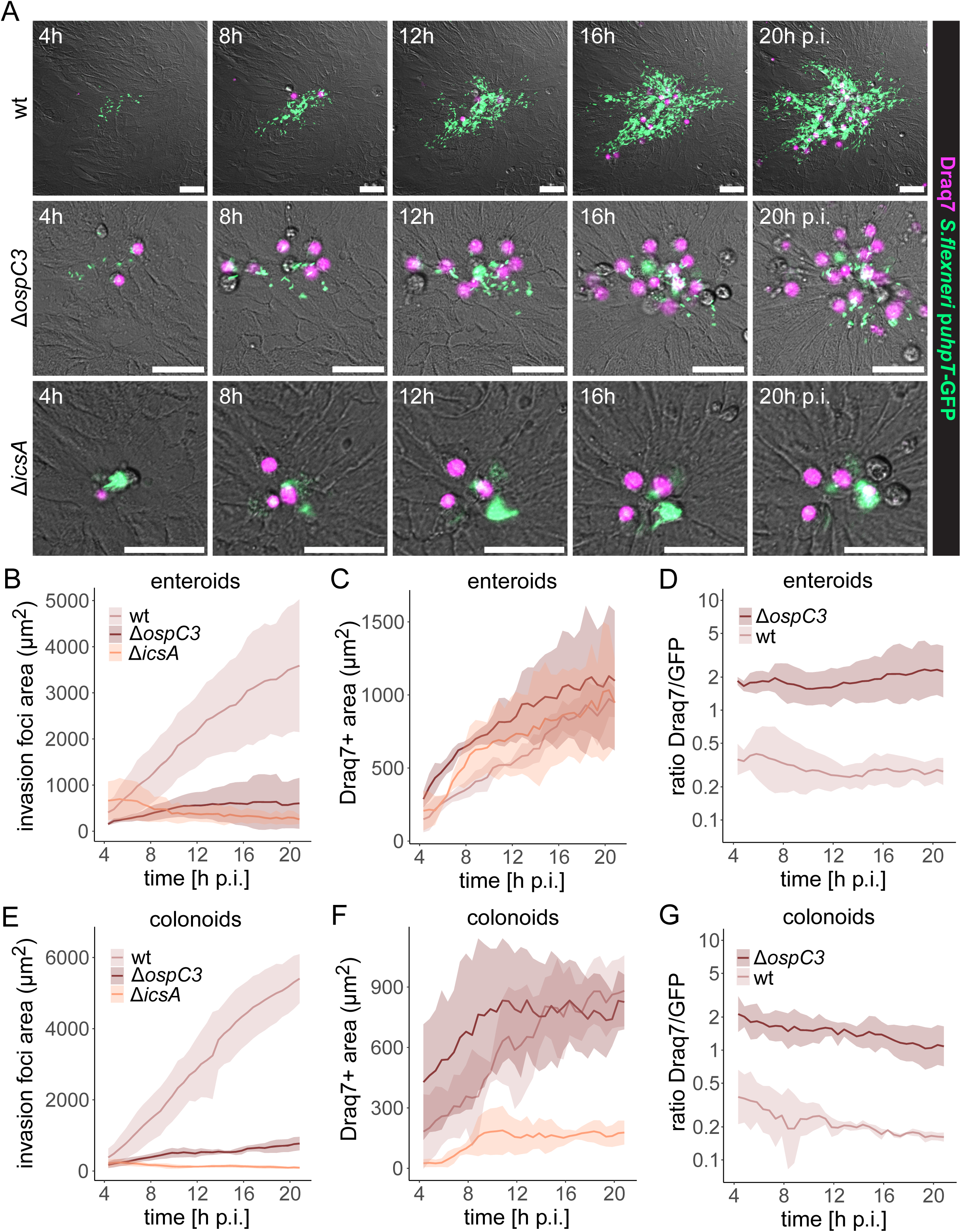
*Shigella* OspC3 suppresses IEC death long enough to enable IcsA-dependent intraepithelial spread. (A-D) Enteroid-derived monolayers were stained with Draq7 and infected with the indicated *Shigella* strains harboring the p*uhpT*-GFP reporter (MOI 200 + centrifugation). Individual invasion foci were followed by time-lapse microscopy and their expansion was quantified. (A) Representative images of *Shigella* wt, but not *Shigella* Δ*ospC3* or *Shigella* Δ*icsA*, expanding intraepithelially. Scale bars: 50µm. Quantification of (B) the GFP-positive area and (C) Draq7-positive area over time. (D) The increased Draq7-to-GFP ratio for *Shigella* Δ*ospC3* indicates a reduced IEC death frequency coupled to successful intraepithelial expansion. Data is plotted as mean + standard deviation (SD) of 3 replicates per strain, with one replicate corresponding to the mean of 3-11 foci in an individually infected well. (E-G) Colonoid-derived monolayers were infected and imaged as in A-D, and (E) GFP-positive area, (F) Draq7-positive area and (G) Draq7-to-GFP ratio were quantified over time. Data is plotted as mean + SD of 3 replicates per strain, with one replicate corresponding to the mean of 2-8 foci in an individually infected well.

To investigate links between epithelial inflammasome-dependent cell death and *Shigella* expansion more generally, we turned to a murine enteroid infection model. While the Caspase-4-dependent non-canonical inflammasome is largely responsible for cell death induction in enterobacterium-infected human intestinal epithelia, the NAIP/NLRC4 inflammasome plays the most prominent role in mouse IECs (45–49, 51). Consequently, WT and *Nlrc4^-/-^* murine enteroid-derived monolayers were infected with *Shigella* wt, incubated with Draq7 and followed by time-lapse microscopy. This revealed the essential absence of stable intraepithelial *Shigella* foci paired with prompt induction of IEC death in WT monolayers, in sharp contrast to lower levels of cell death and efficient *Shigella* foci expansion in *Nlrc4^-/-^* monolayers (Fig S4). These results highlight that a failure of inflammasome-dependent cell death permits rapid lateral *Shigella* expansion also in murine intestinal epithelia.

Finally, when human enteroid/colonoid-derived monolayers were treated with the broad-spectrum caspase inhibitor Z-VAD-FMK, *Shigella* Δ*ospC3* intraepithelial expansion and a correspondingly low Draq7-to-GFP ratio were restored, while the treatment did not further promote *Shigella* wt colonization (Fig 5, enteroids; Fig S5, colonoids). Further consistent with the results above, *Shigella* Δ*icsA* foci failed to expand also under this condition (Fig 5, S5). This demonstrates that inflammasome-dependent, caspase-mediated cell death restricts *Shigella* cell-to-cell spread, but that when cell death is delayed to a certain threshold, either chemically or by a T3SS effector, the intraepithelial *Shigella* population expands rapidly. Altogether, these data highlight that *Shigella* compensates for its poor IEC invasion capacity by the evolution of an efficient intraepithelial expansion module that temporally combines cell death delay by OspC3 and IcsA-mediated cell-to-cell spread to outrun the IEC counter response.

**Figure 5.**
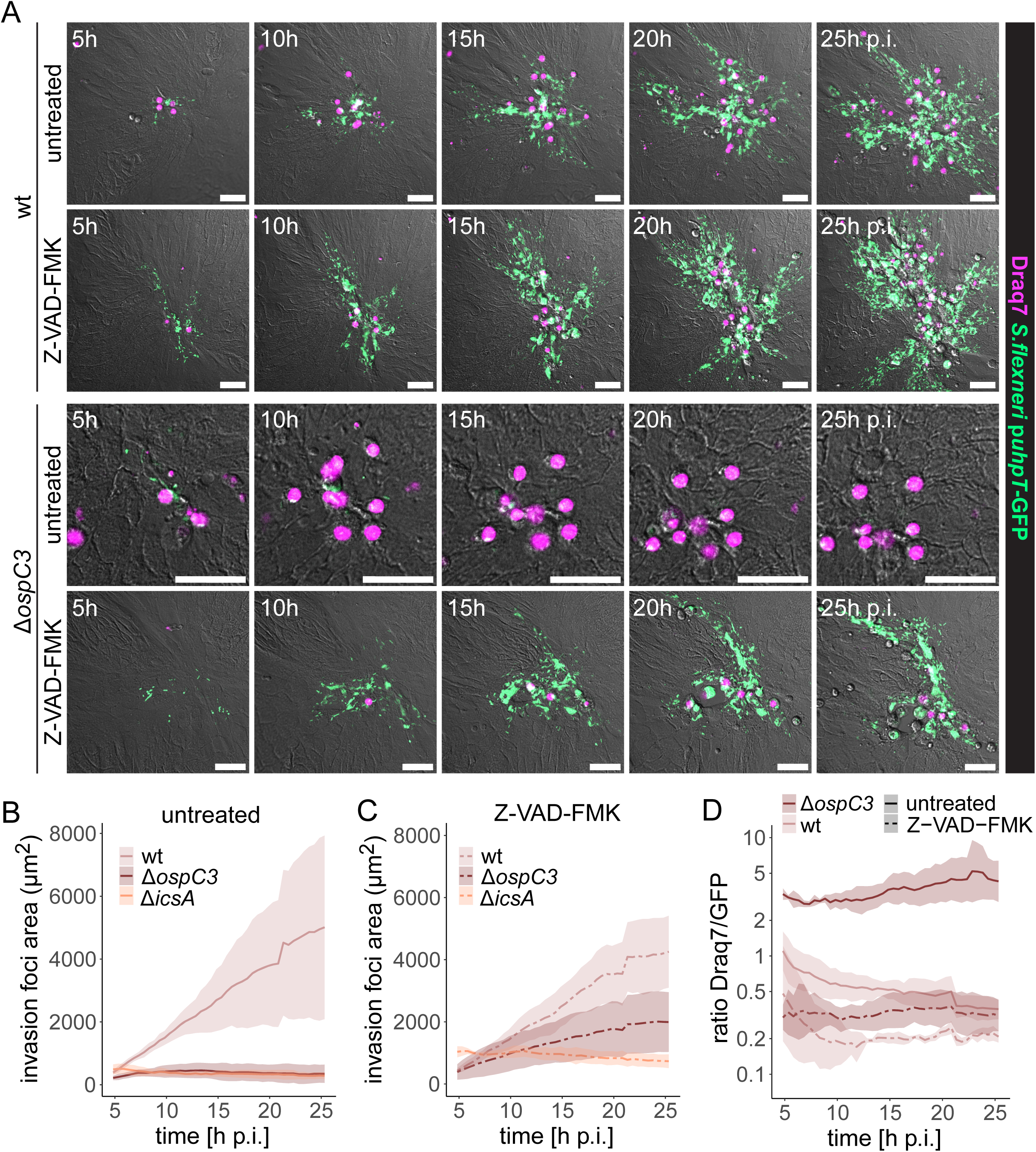
The failure of a *Shigella* Δ*ospC3* mutant to expand intraepithelially can be restored by chemical caspase inhibition. (A-D) Enteroid-derived monolayers were infected with the indicated *Shigella* p*uhpT*-GFP strains as in Figure 4 in the presence or absence of the broad-spectrum caspase inhibitor Z-VAD-FMK and individual invasion foci were followed by time-lapse microscopy. (A) Representative images of Z-VAD-FMK treatment restoring *Shigella* Δ*ospC3* intraepithelial expansion to wt levels. Scale bars: 50µm. (B-C) Quantification of the GFP-positive area in (B) untreated and (C) Z-VAD-FMK-treated monolayers over time. (D) The Draq7-to-GFP ratio for *Shigella* Δ*ospC3* was reduced to wt levels upon Z-VAD-FMK treatment. Data is plotted as mean + SD of 3 replicates per strain and treatment, with one replicate corresponding to the mean of 3-10 (untreated) or 5-12 (Z-VAD-FMK) foci in an individually infected well.

## DISCUSSION

Previous work on enterobacterial virulence factors and epithelial host cell responses has provided a rich catalogue of infection-relevant biochemical interactions. These studies have placed strong emphasis on molecular mechanism. However, less attention has been paid to recapitulating primary cell behavior and resolving temporal interconnections between virulence factor sets and host cell responses throughout infection stages. By employing time-lapse imaging in human enteroids and colonoids that retain primary IEC features and functional cell death responses, we here generated a dynamic, spatially and temporally resolved map of the divergent *Salmonella* and *Shigella* epithelial colonization strategies.

Equipped with an apical targeting module combining flagellar momentum and adhesin activity, *Salmonella* efficiently scans the surface of the intestinal epithelium and encounters a high density of suitable binding sites. The SPI4-encoded SiiE adhesin system, which we identified as central for *Salmonella* attachment to the apical enteroid/colonoid surface, has previously been attributed an adhesive role for invasion of polarized cell lines and *in vivo* in mice (14, 29). Here we extend these findings by using uniquely designed adhesion assays to reveal how SiiE enables transition from Brownian-like floating atop the epithelial surface to stable adhesion, likely accomplished through multivalent binding to glycosylated membrane-bound mucins like MUC1 and MUC13 (68, 69). This step preceded and enables T3SS-1-driven invasion. *Salmonella* expresses a range of other adhesins under various conditions (10–13), so the prominent role of SiiE identified here could be replaced by other adhesins in other host species. Notably, the importance of SPI4-dependent binding appears consistent across our experiments in human enteroids and colonoids that derive from vastly different gut segments. While flagella have also been ascribed adhesive properties (8, 9), our assays provided no evidence for such function in apical binding to non-transformed human IECs. Rather, flagellar motility allowed *Salmonella* to partially overcome the requirement for adhesins, likely by generating momentum to penetrate the apical brush border and stably dock to the IEC membrane via T3SS-1. Collectively, the collaboration of three virulence systems, flagella, a SPI-4-encoded adhesin and T3SS-1, mediates *Salmonella* targeting, stable docking and apical IEC invasion at reasonable frequency in human enteroids/colonoids. This promptly induces cell death and highlights the epithelial inflammasome response as the most prominent restrictive mechanism for long-term *Salmonella* epithelial colonization (Fig S6).

Removal of the apical targeting module (*ΔfljBfliC ΔSPI-4*) effectively transformed *Salmonella* into a *Shigella*-like particle stalled floating atop the IEC surface. This highlights the lack of well-described adhesins and flagella as the decisive differences between *Shigella* and *Salmonella* during early IEC invasion, and suggests that cooperation with external factors are needed for *Shigella* to establish an initial foothold in human absorptive epithelia. Indeed, this notion is supported by other *in vitro* models for *Shigella* infection relying on artificial adhesin expression or a centrifugation step to enforce host cell contact (70, 71). The requirement for preexisting epithelial damage to colonize the enteroid/colonoid epithelium from the apical side in our microinjection model, supposedly by exposing the lateral or basal membrane for T3SS docking, offers a plausible mechanism of how this barrier might be occasionally breached *in vivo*. Transit through M-cells has been described as an entry portal for *Shigella* (24, 25, 28, 60, 72), but studies in other models have also shown that infiltrating polymorphonuclear leukocytes (28, 59), malnourishment (61) or EGTA treatment (22, 23), increase epithelial permeability and may allow for *Shigella* invasion analogous to what we find for physical damage. Furthermore, host factors have been suggested to directly (20, 27) or indirectly (17–19) promote *Shigella* adhesion. Nevertheless, *Shigella* apical invasion of non-transformed human absorptive epithelia by own efforts appears exceedingly rare, highlighting the IEC binding/invasion step as a major barrier for *Shigella* infection, and explaining the critical need for cell death silencing and expansion mechanisms once this barrier is occasionally overcome (Fig S6).

At the post-invasion step, we indeed found that the concerted action of the inflammasome antagonist OspC3 and the actin nucleator IcsA allowed for lateral spread just in time to avoid clearing by IEC death. In the absence of OspC3, *Shigella* invasion foci were frequently cleared, but caspase inhibition by chemical means again allowed the pathogen to laterally outrun the epithelial cell death response and successfully maintain an infection focus. While the non-canonical Caspase-4 pathway targeted by OspC3 represents the main inflammasome response in human IECs (45, 47, 51), the NAIP/NLRC4-Caspase-1 pathway most prominent in the murine epithelium (45, 46, 48) is unaffected by *Shigella* effectors. This dichotomy in epithelial inflammasomes prevents *Shigella* colonization of the murine gut epithelium, as shown in a recent study (44), and further temporally explained by our live-cell imaging experiments in murine enteroid-derived monolayers.

Collectively, we describe two virulence factor modules giving rise to divergent strategies for successful colonization of human absorptive intestinal epithelia by prominent representatives of invasive enterobacteria. The *Salmonella* “polyclonal and iterative” colonization strategy relies on the apical targeting module and is vastly superior in all steps prior to entry. This results in multiple independent, but relatively short-lived IEC invasion foci, and exploits inflammasome-mediated IEC extrusion for luminal re-seeding and expansion of a bacterial population primed for reinvasion (57, 73, 74) (Fig S6). *Shigella*, on the other hand utilizes an “essentially monoclonal” colonization strategy marked by exceedingly rare IEC invasion events paired with superiority in steps subsequent to entry. Here, the identified intraepithelial expansion module comprised of OspC3 and IcsA pairs temporal delay of the epithelial inflammasome response (and potentially other pro-inflammatory and cell death pathways) with exploiting the host cell actin machinery for cell-to-cell spread just in time before a colonized IEC succumbs (Fig S6). This remarkable expansion capacity in non-transformed epithelia may also explain the sufficiency of the low *Shigella* infectious dose described in clinical studies (1). It is moreover noteworthy that the sensitive cell death responses of primary IECs do not permit the expansive intracellular replication seen in cell lines, but the *Shigella* strategy still enables sufficient rounds of intra-IEC replication for plaque expansion by the progeny of a single founder bacterium. Finally, considering host range, it is tempting to speculate that host-restricted epithelial invaders such as *Shigella* are more likely to adapt a recognition-suppressive strategy subsequent to IEC entry, while generalist invaders such as *Salmonella* would favor high efficiency at the entry step, due to the infeasibility of suppressing immune recognition pathways that can differ markedly across host species.

Despite primary IEC features and cell death responses in enteroids/colonoids, the lack of a fully established soluble mucus layer, and the absence of microbiota and immune cells, should be pointed out among the limitations of this study. However, enterobacterial pathogens oftentimes target sites devoid of a continuous mucus layer, such as the murine cecum (6), or the follicle-associated epithelium featuring a penetrable mucus layer (75). In addition, more advanced organoid models incorporating one or several of the abovementioned aspects are being developed (59, 76–79), and will be valuable for future studies building on this one. The current setup nevertheless allows to follow *Salmonella* and *Shigella* epithelial colonization in non-transformed human epithelia in real-time, revealing how modules of bacterial species-specific virulence factors active at the apical targeting step (*Salmonella*) versus the intracellular expansion step (*Shigella*) dictate vastly different epithelial colonization dynamics (Fig S6).

## MATERIALS AND METHODS

### Ethics statement

Human enteroids were established from jejunal tissue resected during routine bariatric surgery. Human colonoids were established from morphologically normal non-tumor tissue resected during elective colon cancer surgery. All tissue samples were acquired following each subject’s informed consent. Personal data were pseudonymized prior to further processing in the lab, such that patient identities were unknown to the researchers working on the samples. All procedures for establishment and experimentation on enteroids and colonoids were approved by the local governing body (Etikprövningsmyndigheten, Sweden) under license numbers 2010-157 (with addenda 2010-157-1 and 2020–05754) and 2023-01524-01. Optimization of protocols for enteroid and colonoid establishment, 2D and 3D growth, and infection experiments, involved several independent enteroid/ colonoid cultures. Final data sets presented in figures derive from cultures Hu18-9jej and Hu22-4col.

### Bacterial strains, plasmids and culture conditions

All bacterial strains and reporter plasmids used in this study are listed in Table S1. *Salmonella enterica* serovar Typhimurium (*Salmonella*, *S*.Tm) strains were of SL1344 background (SB300, streptomycin resistant, Sm^R^) (80). Besides the wild-type (wt), previously described Δ*invG*, Δ*SPI-4* and Δ*fljBfliC* mutants were used. The Δ*fljBfliC* Δ*invG*, Δ*fljBfliC* Δ*SPI-4* and Δ*motA* Δ*SPI-4* mutants were generated by flip-out of the kanamycin resistance cassette (Kan^R^) from the aforementioned Δ*fljBfliC* and previously published Δ*motA* (57) strains, followed by transfer of previously described deletions from *Salmonella* 14028 (C1793) (81) or SL1344 strains (Δ*SPI-4*) (14), respectively, by P22 transduction. Genotypes were verified by PCR using primer pairs k1/invG del scr F, k2/invG del scr R and invG del scr F/invG del scr R for Δ*fljBfliC* Δ*invG* and primer pairs k1/SPI4-Ctrl-For, k2/SPI4-Ctrl-Rev and SPI4-Ctrl-For/SPI4-Ctrl-Rev for Δ*fljBfliC* Δ*SPI-4* and Δ*motA* Δ*SPI-4* (Table S2). All *Shigella flexneri* (*Shigella*, *S.fl*) strains were of M90T background (82). Besides the wt, the previously described Δ*mxiD* mutant was used. Δ*ospC3* and Δ*icsA* mutants were generated by lambda red recombination (83), transforming M90T pSIM5 (84) with the PCR product obtained using plasmid pKD13 as template and the primer pairs ospC3-Del-fwd/ospC3-Del-rev and icsA-Del-fwd/icsA-Del-rev, respectively (Table S2). Genotypes were verified by PCR using the primer pairs ospC3-Ctrl-fwd/ ospC3-Ctrl-rev and icsA-Ctrl-fwd/ icsA-Ctrl-rev, respectively (Table S2). The constitutive pFPV-mCherry (*rpsM*-mCherry; Addgene plasmid number 20956) (65) and intracellular pM975 (p*ssaG*-GFPmut2) (48, 63) and pZ1400 (p*uhpT*-GFP) (64) reporter plasmids (Table S1) were described and validated previously. For infections, *Salmonella* strains were grown overnight at 37°C for 12 h in LB/0.3 M NaCl (Sigma-Aldrich) with appropriate antibiotics and sub-cultured (1:20 dilution) in the same medium without antibiotics at 37°C for 4 h. *Shigella* strains were grown overnight at 30°C for 16 h in LB with appropriate antibiotics and sub-cultured (1:50 dilution) in LB without antibiotics at 37°C for 2 h until OD600 ∼0.7. For Caco-2 infections, the inoculum was reconstituted in DMEM GlutaMAX (Gibco)/0.1 mM non-essential amino acids (NEAA) (Gibco) at a concentration of 2x 10^7^ CFU/ml, while for enteroid/colonoid infections, it was reconstituted in antibiotic-free human (hOGM) or mouse (mOGM) IntestiCult organoid growth medium (StemCell) at a concentration of ∼10^9^ CFU/ml and further diluted 1:100 for low MOIs where indicated.

### Human enteroid and colonoid establishment

Human jejunal enteroids and colonoids were established as previously described (57, 85). Newly established enteroids and colonoids were frozen at passage 1 to 4 by gently dissolving Matrigel (Corning) domes in ice-cold DMEM/F12 (Gibco)/0.25% BSA (Gibco) and resuspending the enteroids/colonoids in DMEM/F12/10% heat-inactivated fetal bovine serum (FBS) (Gibco)/10% DMSO (Sigma-Aldrich), followed by freezing in a Mr. Frosty^TM^ Freezing container at −80°C overnight and transfer to a liquid nitrogen tank for cryopreservation.

### Human and murine enteroid/colonoid culture

Human jejunal enteroids and colonoids were cultured in Matrigel (75%) domes overlaid with hOGM/1x PenStrep (Gibco) at 37°C, 5% CO_2_ and passaged once a week as previously described (67). Enteroids from passages 4 to 30, and colonoids from passage 3 to 16 were used for experimentation. WT (86) and *Nlrc4-/-* (50, 87) murine jejunal enteroids of C57BL/6 background were cultured in Matrigel (60%) domes overlaid with mOGM/1x PenStrep at 37°C, 5% CO_2_ and passaged every 5-7 days as previously described (67) and passages 10 to 16 were used for experimentation.

### Caco-2 cell culture and infection

Caco-2 cells were cultured in DMEM/10% heat-inactivated FBS/0.1 mM NEAA/1x PenStrep at 37°C, 10% CO_2_ and passaged every 2 to 3 days. 1 day prior to infection, cells were seeded at ∼55’000 cells/cm^2^ in a glass-bottom 24-well plate (Cellvis) pre-coated with 100 µg/cm^2^ collagen I (Corning). For infection, the medium was exchanged for half the volume of the well of antibiotic-free DMEM/0.1 mM NEAA, and wells were filled by adding the same volume of the prepared inoculum. The plate was centrifuged at 700 g for 10min, incubated at 37°C, 5% CO_2_ for 40 min and washed 3 times with DMEM/10% FBS/0.1 mM NEAA before addition of DMEM/10% FBS/0.1 mM NEAA/200 µg/ml gentamicin (Sigma-Aldrich).

### Human enteroid/colonoid microinjection

Microinjection of *Salmonella* and *Shigella* into human enteroids and colonoids was performed as previously described (57). Briefly, enteroids/colonoids were passaged, embedded in Matrigel (∼100%) in 35-mm glass-bottom dishes (MatTek) and overlaid with 2 ml antibiotic-free hOGM. The culture medium was replaced every 3 days and prior to microinjection. Enteroids were injected 5 to 7 days after passaging, while colonoids were injected 7 to 8 days after passaging due to the slightly slower growth rate. Microinjection needles were prepared from 1.0-mm filamented glass capillaries (World Precision Instruments; no. BF100-78-10; Borosilicate, 1 mm wide, 100 mm long, with filament) using a micropipette puller (Sutter Instruments; P-1000) and the following settings: heat = ramp + 5; pull = 60; velocity = 80; delay = 110; pressure = 200. Needles were beveled at a 30° angle on a fine-grit diamond lapping wheel, loaded with the *Salmonella* or *Shigella* inocula by fluidic force and mounted on a microinjector (MINJ-FLY; Tritech Research) controlled by a micromanipulator (uMP-4; Senapex). 20-200 ms pressured air pulses were applied to inject ∼50-500 bacteria, depending on the size of the enteroid/colonoid.

### Establishment and infection of 2D human enteroid/colonoid-derived monolayers

Human enteroid and colonoid-derived monolayers were established on Matrigel-coated polymer coverslips (µ-Slide 8-well high Cat.No 80806 or µ-Plate 96-well black Cat.No 89626, Ibidi) as previously described (67). Briefly, enteroids and colonoids were used for monolayer establishment 7 days after passaging. Enteroids/colonoids were extracted from Matrigel domes using Gentle Cell Dissociation Reagent (StemCell), washed once in DMEM/F12/1.5% BSA and dissociated into single cells using TrypLE (Gibco) and mechanical shearing by pipetting. The single cell suspension was washed one more time in DMEM/F12/1.5% BSA, reconstituted in antibiotic-free hOGM/10 µM Y-27632 (Sigma-Aldrich) and seeded out at 750’000 cells/cm^2^ in wells pre-coated with a 1:40 Matrigel dilution in DPBS (Gibco). Monolayers were maintained at 37°C, 5% CO_2_ and 2-3 days post establishment, the medium was exchanged for fresh antibiotic-free hOGM without Y-27632. Monolayers were used for infection 3 to 5 days post establishment. After washing with DMEM/F12, half the volume of the well of antibiotic-free hOGM, supplemented with 1.5 µM Draq7 (Invitrogen) and/or 100 µM Z-VAD-FMK (AH diagnostics) where indicated, was added and the wells were filled by adding the same volume of the prepared inoculum. Unless indicated otherwise, slides were centrifuged at 700 g for 10 min prior to incubation at 37°C, 5% CO_2_ for 40 min to allow for bacterial invasion. After washing 3 times with DMEM/F12/400 µg/ml gentamicin, hOGM/50 µg/ml gentamicin, supplemented with 0.75 µM Draq7 and/or 50 µM Z-VAD-FMK where indicated, was added to the wells.

### Establishment and infection of 2D murine enteroid-derived monolayers

For monolayer establishment, murine enteroids were pre-treated with mOGM/1 mM valproic acid (Cayman chemicals)/3 µM CHIR99021 (Cayman chemicals) (CV medium) for 7 days as previously described (67, 85). Enteroids were extracted from Matrigel domes using Gentle Cell Dissociation Reagent, washed once in DMEM/F12/1.5% BSA and dissociated into single cells using TrypLE and mechanical shearing by pipetting. The single cell suspension was washed one more time in DMEM/F12/1.5% BSA, reconstituted in antibiotic-free CV medium/10 µM Y-27632 and seeded out at 530’000 cells/cm^2^ in wells pre-coated with a 1:40 Matrigel dilution in DPBS. Monolayers were maintained at 37°C, 5% CO_2_ and 24-36 h post establishment, the medium was exchanged for fresh antibiotic-free mOGM without Y-27632. Monolayers were used for infection 3 days post establishment. After washing with DMEM/F12, half the volume of the well of antibiotic-free mOGM supplemented with 1.5 µM Draq7 was added and the wells were filled by adding the same volume of the prepared inoculum. Slides were centrifuged at 700 g for 10 min, incubated at 37°C, 5% CO_2_ for 40 min and washed 3 times with DMEM/F12/400 µg/ml gentamicin, followed by addition of mOGM/50 µg/ml gentamicin.

### Fixation and staining of enteroid/colonoid-derived monolayers

3-4 h p.i., enteroid and colonoid-derived monolayers were washed twice with DPBS and fixed with DPBS/2% paraformaldehyde for 15 min in the dark. Following two additional washes with DBPS, they were stained with 1 µg/ml DAPI (Sigma-Aldrich) and 1 U/ml Phalloidin-Alexa647 (Molecular probes) in DPBS/0.1% Triton X-100 (Sigma-Aldrich) for 30 min, shaking in the dark. After two final washes as described above, DPBS was added to the wells prior to image acquisition.

### Microscopy

Imaging was performed on a custom-built microscope based on an Eclipse Ti2 body (Nikon), using 60x/0.7 and 40x/0.6 Plan Apo Lambda air objectives (Nikon) and a back-lit sCMOS (scientific complementary metal oxide semiconductor) camera with pixel size 11 µm (Prime 95B; Photometrics). For time-lapse microscopy, the imaging chamber was maintained at 37°C, 5% CO_2_ in a moisturized atmosphere. Bright-field images were acquired by differential interference contrast (DIC), and fluorescence imaging by the excitation light engine Spectra-X (Lumencor) and emission collection through a quadruple band pass filter (89402; Chroma). Infected Caco-2 cells were imaged every 2 min for 16 h. Microinjected enteroids and colonoids were imaged immediately after microinjection, and time-lapse images were acquired every 5-15 min for up to 16 h. For bacterial tracking, microinjected enteroids and colonoids were imaged at 500-ms intervals for 20 frames in total at each indicated time point. Z-stacks of non-centrally injected enteroids and colonoids were acquired between the middle and bottom plane at the indicated time points with 2-5 µm between slices. Manually detected invasion foci in enteroid/colonoid-derived monolayers were imaged every 30 min for up to 25 h. Automated image acquisition of fixed enteroid-derived monolayers was performed using Micromanager 2.0 (88), while fixed colonoid-derived monolayers were imaged manually to include rare and unevenly distributed invasion foci.

### Image analysis

Background subtraction for fluorescence channels was performed in Fiji (a version of ImageJ) (89) applying a rolling ball radius of 50 pixels. The TrackMate plugin (90) in Fiji was used for tracking and all tracks were verified manually due to bacterial crowding at the bottom plane complicating automated tracking. For quantification of the fluorescence retained at the side epithelium of microinjected enteroids, maximum intensity projections of background-subtracted z-stacks from the mCherry channel were generated in Fiji and middle and bottom plane cross sections were delineated based on DIC images. The same threshold was set for all time points from one replicate and the fluorescence above threshold for the non-overlapping area of the middle and bottom plane cross sections (i.e. the side epithelium) was quantified in Fiji. A CellProfiler (91) customized pipeline was used for quantification of invasion foci in fixed enteroid/colonoid-derived monolayers. For quantification of invasion foci and Draq7-positive areas, one threshold value per channel was used for all time-lapse movies from one experiment, and the area above threshold was quantified in Fiji.

### Statistical analysis

All data was analyzed and plotted using the tidyverse collection of R packages (92) in RStudio (93).

## ACKNOWLEDGEMENTS

We are grateful to members of the Sellin laboratory for helpful discussions and to the staff of our surgical units for assistance with human intestinal tissue sampling. This work was supported by grants from the Swedish Research Council (2018-02223, 2022-01590), the Swedish Foundation for Strategic Research (FFL18-0165), and the SciLifeLab Fellows program to M.E.S.

## DISCLOSURE OF INTEREST

The authors declare no competing interests.

## AUTHOR CONTRIBUTIONS

Conceptualization: P.G., M.E.S.; methodology: P.G., M.L.D.M., A.C.C.L., A.B., J.E.; investigation: P.G., M.L.D.M., A.C.C.L.; formal analysis: P.G.; interpretation: P.G., M.L.D.M, M.E.S.; resources: M.L.D.M., A.L., M.Su., M.Sk., W.G., D.-L.W., P.M.H.; supervision: M.L.D.M., M.Su., M.E.S.; project administration: M.Su., M.Sk., W.G., D.-L.W., P.M.H., M.E.S.; funding acquisition: M.E.S., visualization: P.G., writing – original draft: P.G., M.E.S.; writing – reviewing and editing: all authors.

## Supplementary Material for

**Supplementary Figure S1.**
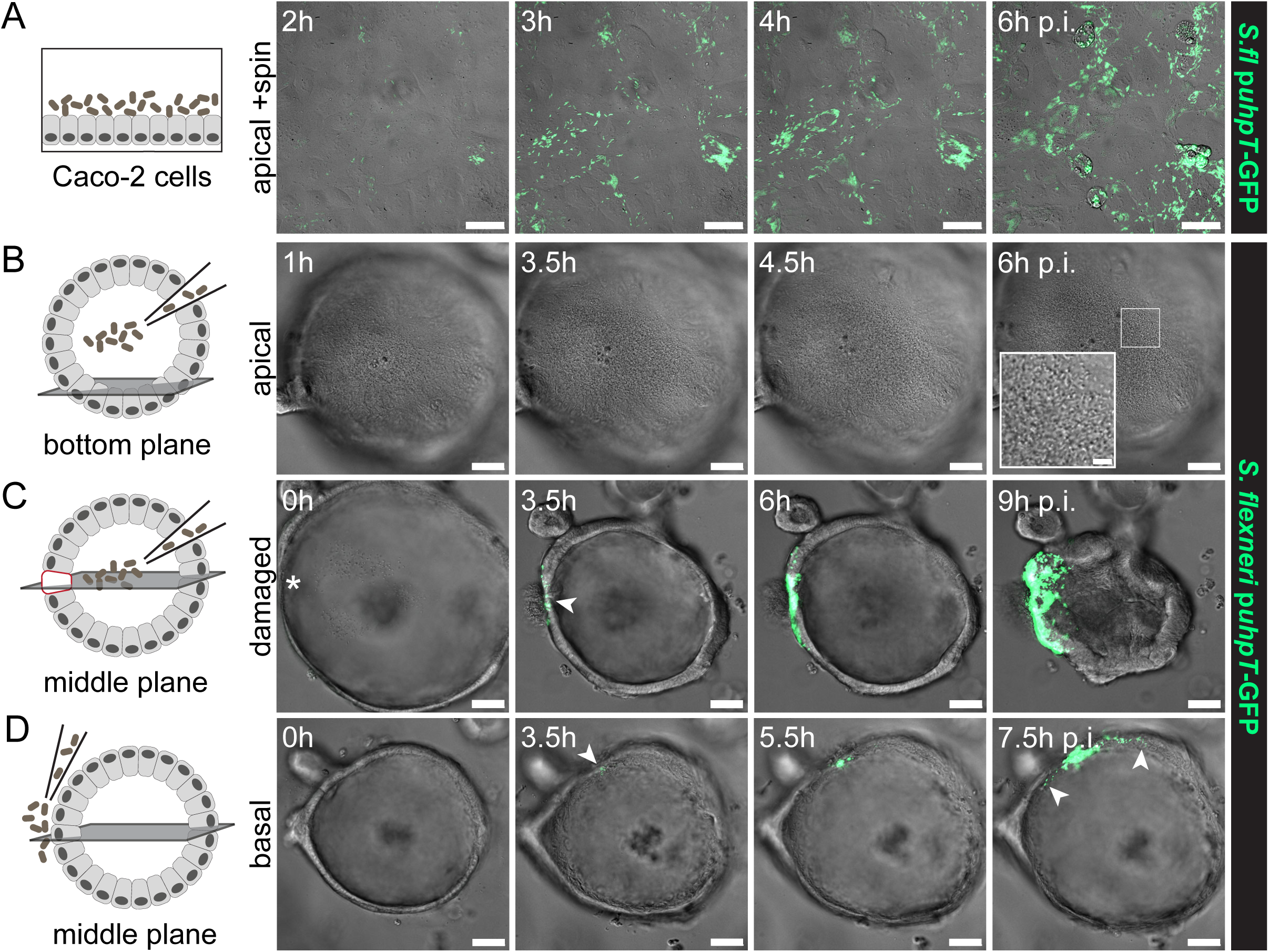
Epithelial damage results in localized *Shigella* invasion of the colonoid epithelium from the apical side. (A) Caco-2 cells were apically infected with *Shigella* wt p*uhpT*-GFP (MOI 140), including a centrifugation step, and imaged by DIC and fluorescence time-lapse microscopy. (B-D) Human colonoids were infected with *Shigella* wt p*uhpT*-GFP via different routes and imaged as in A. (B) Upon luminal microinjection, *Shigella* quickly accumulates at the bottom plane (see insert), but the unperturbed epithelium was not permissive for invasion from the apical side. (C) Epithelial damage introduced with the microinjection needle (*) resulted in localized invasion from the site of damage. (D) Basolateral deposition of *Shigella* also resulted in successful epithelial colonization. Arrowheads indicate invasion foci. Scale bars: 50µm (10µm for insert). *S.fl*, *Shigella flexneri*.

**Supplementary Figure S2.**
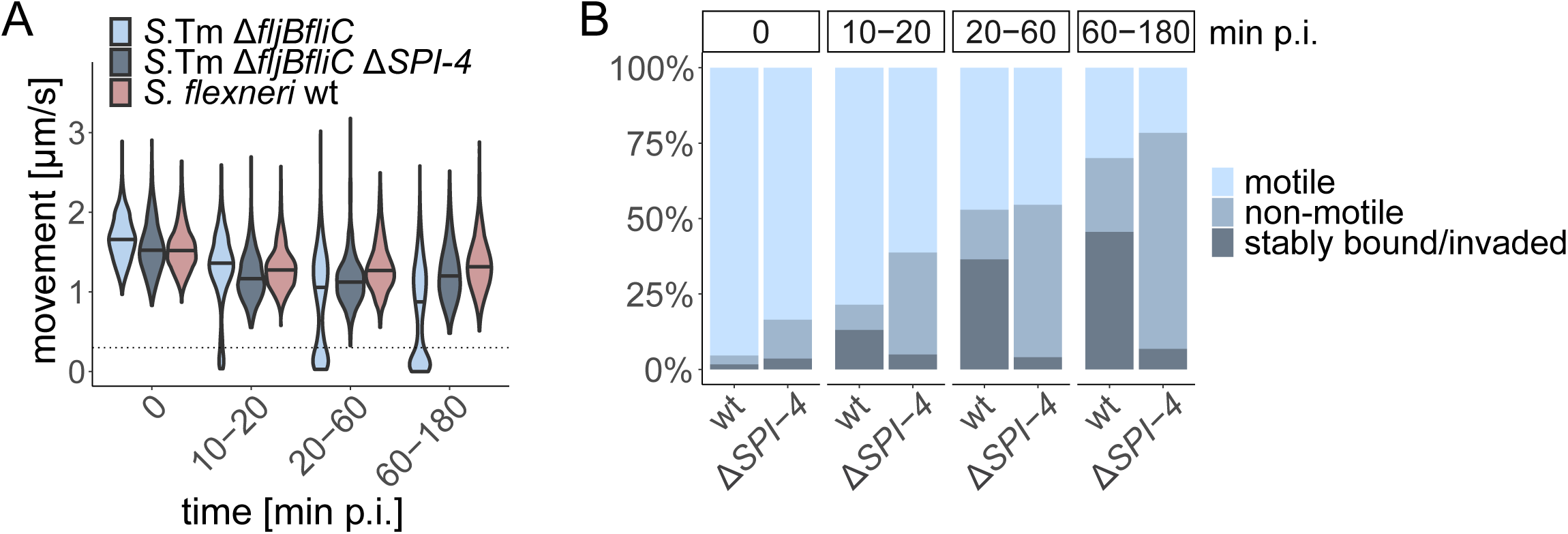
The SiiE adhesin and flagellar motility promote adhesion to the enteroid and colonoid epithelium. (A) Colonoids were microinjected with the indicated *Salmonella* and *Shigella* strains harboring the constitutive *rpsM*-mCherry reporter, and bacterial movement within the colonoids was tracked. The movement speeds at the at the indicated time points p.i. were quantified. The bimodal population distribution for *Salmonella* Δ*fljBfliC* with movement speeds close to 0 indicates a stably adhering population. Data from at least 9 colonoids and at least 430 tracks per time point and strain. Horizontal lines indicate the median. (B) Enteroids were microinjected with *Salmonella* wt and *Salmonella* Δ*SPI-4 rpsM*-mCherry and bacterial movement within the enteroids was tracked. Tracks were classified as motile (>5µm/s), non-motile (0.3-5µm/s) or stably bound/invaded (<0.3µm/s) based on their swimming speeds and plotted as percentage of the total population. Data from at least 7 enteroids and at least 310 tracks per time point and strain.

**Supplementary Figure S3.**
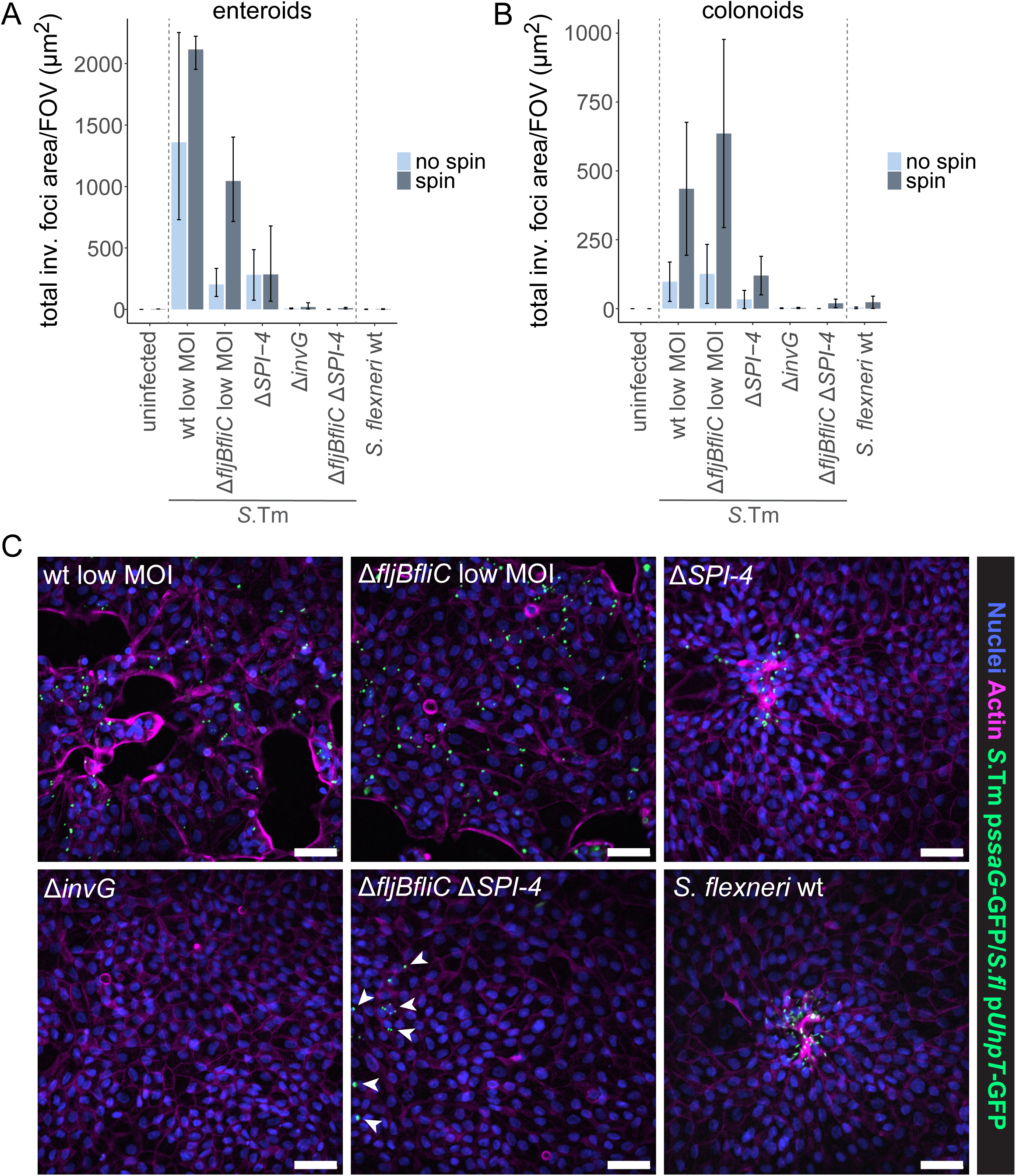
The SiiE adhesin and flagellar motility both promote invasion of the enteroid and colonoid epithelium. (A) Enteroid-derived monolayers were infected with the indicated *Salmonella* and *Shigella* strains at MOI 200, or MOI 2-4 (“low MOI”), respectively, for 40min, with or without centrifugation. Monolayers were fixed at 3-4 h p.i. and the total area of invasion foci at randomly selected fields of view (FOVs) was determined based on the intracellular reporters p*ssaG*-GFP and p*uhpT*-GFP, respectively. Data is plotted as mean + range of 3 replicates per strain (1 replicate for uninfected control), with one replicate corresponding to the mean of 14-16 FOVs for an individually infected well. (B-C) Colonoid-derived monolayers were infected as in A, but FOVs for analysis were selected manually to also include rare and unevenly distributed invasion foci. (B) Data is plotted as mean + range of 2 replicates per strain (1 replicate for uninfected control without spin), with one replicate corresponding to the mean of 3-15 (no spin) or 6-17 (spin) FOVs for an individually infected well. (C) Representative images of infected colonoid-derived monolayers (+ centrifugation) stained with DAPI (nuclei) and phalloidin (actin). Scale bars: 50µm. *S*.Tm, *Salmonella* Typhimurium; *S.fl*, *Shigella flexneri*.

**Interpretation.** Enteroid-derived monolayers were infected with different *Salmonella* and *Shigella* strains with or without a centrifugation step to force contact of the bacteria with the epithelial surface in order to determine the requirement for flagellar motility and the SPI-4 adhesin system for invasion under both conditions. The infections were allowed to proceed for 40 min such that even the non-motile bacteria could reach the surface of the monolayers by gravity in the absence of centrifugation. Adhesin-proficient *Salmonella* wt and *Salmonella* Δ*fljBfliC* invaded monolayers with high efficiency that further increased upon centrifugation, particularly for the non-flagellated strain (Fig S3A). Due to the observed reduction in bacterial binding, infections with adhesin-deficient *Salmonella* strains and *Shigella* were performed at 50-100x higher MOI. While invasion by *Salmonella ΔSPI-4* was lower than for the wt even at increased MOI, it was completely undetectable for *Salmonella* Δ*fljBfliC ΔSPI-4* and *Shigella* wt regardless of centrifugation (Fig S3A). This confirms previous results and highlights the importance of the *Salmonella* SPI-4 adhesin system not only for successful adhesion to, but also subsequent invasion of the enteroid epithelium, and that this requirement for adhesins can be partially circumvented by flagellar motility. When repeating the experiment in colonoid-derived monolayers and manually choosing the FOVs for analysis, it was observed that, upon centrifugation, even *Salmonella* Δ*fljBfliC ΔSPI-4* and *Shigella* wt were occasionally able to invade the epithelium, but these invasion foci were exceedingly rare and unevenly distributed with hotspots for invasion determined by monolayer topology (Fig S3B-C). Furthermore, it was found that although the invasion efficiency for *Salmonella ΔSPI-4* was higher than for its non-flagellated counterpart, invasion foci displayed a similarly uneven distribution (Fig S3C). Adhesion-competent *Salmonella* wt and *Salmonella* Δ*fljBfliC*, on the other hand, colonized the colonoid epithelium more evenly and efficiently, thereby also inducing significant levels of IEC death, as revealed by holes in the monolayers (Fig S3C).

**Supplementary Figure S4.**
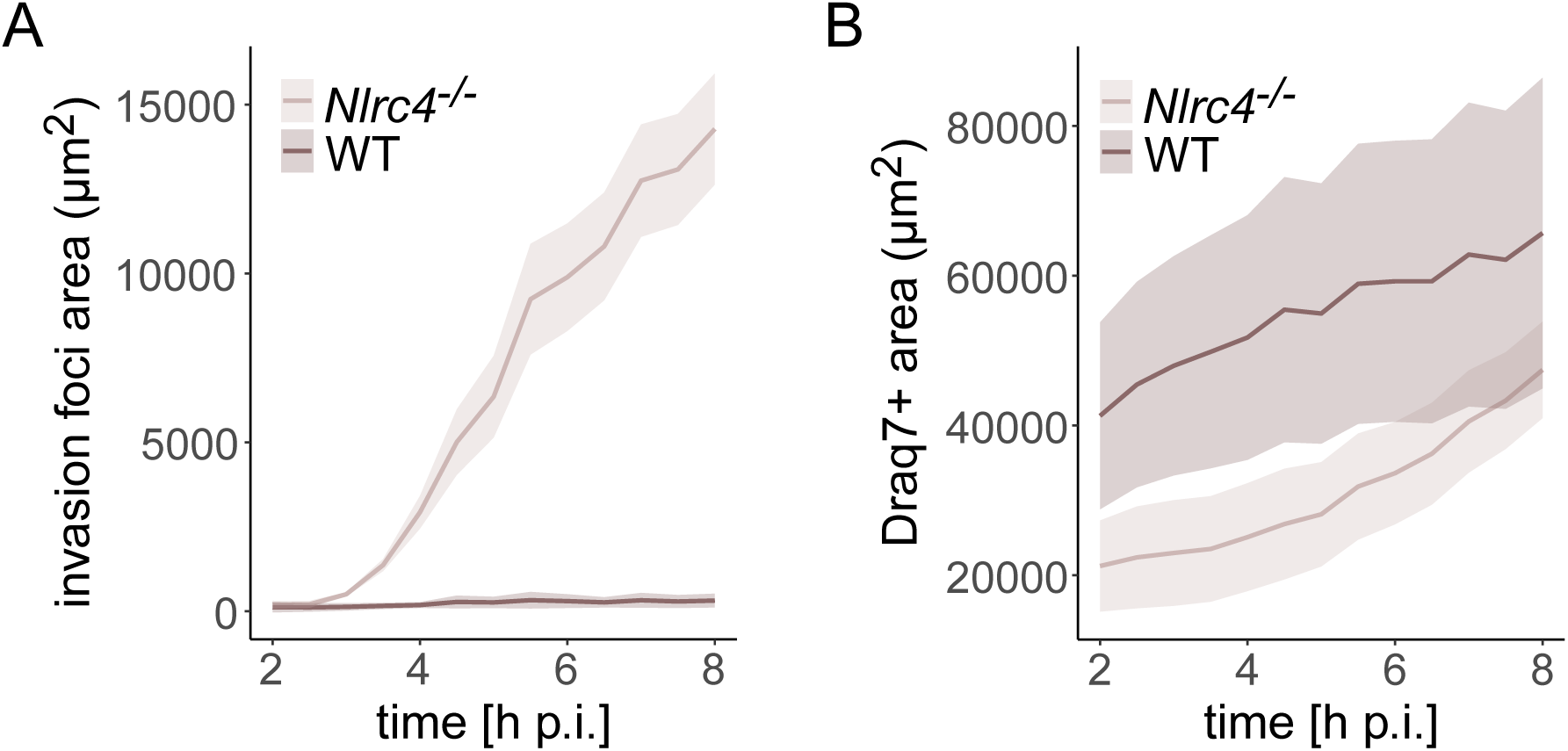
*Nlrc4*-deficient, but not WT murine enteroid-derived monolayers are permissive for *Shigella* colonization. (A-B) WT and *Nlrc4^-/-^* murine enteroid-derived monolayers were stained with Draq7 and infected with *Shigella* wt p*uhpT*-GFP (MOI 200 + centrifugation). (A) Quantification of the GFP-positive area indicates that only *Nlrc4^-/-^*monolayers are permissive for *Shigella* invasion and subsequent intraepithelial expansion. (B) Quantification of the Draq7-positive area reveals reduced cell death in infected *Nlrc4^-/-^* monolayers. Data is plotted as mean + SD of 3-4 replicates per genotype, with one replicate corresponding to the mean of 3-4 fields of view for an individually infected well.

**Supplementary Figure S5.**
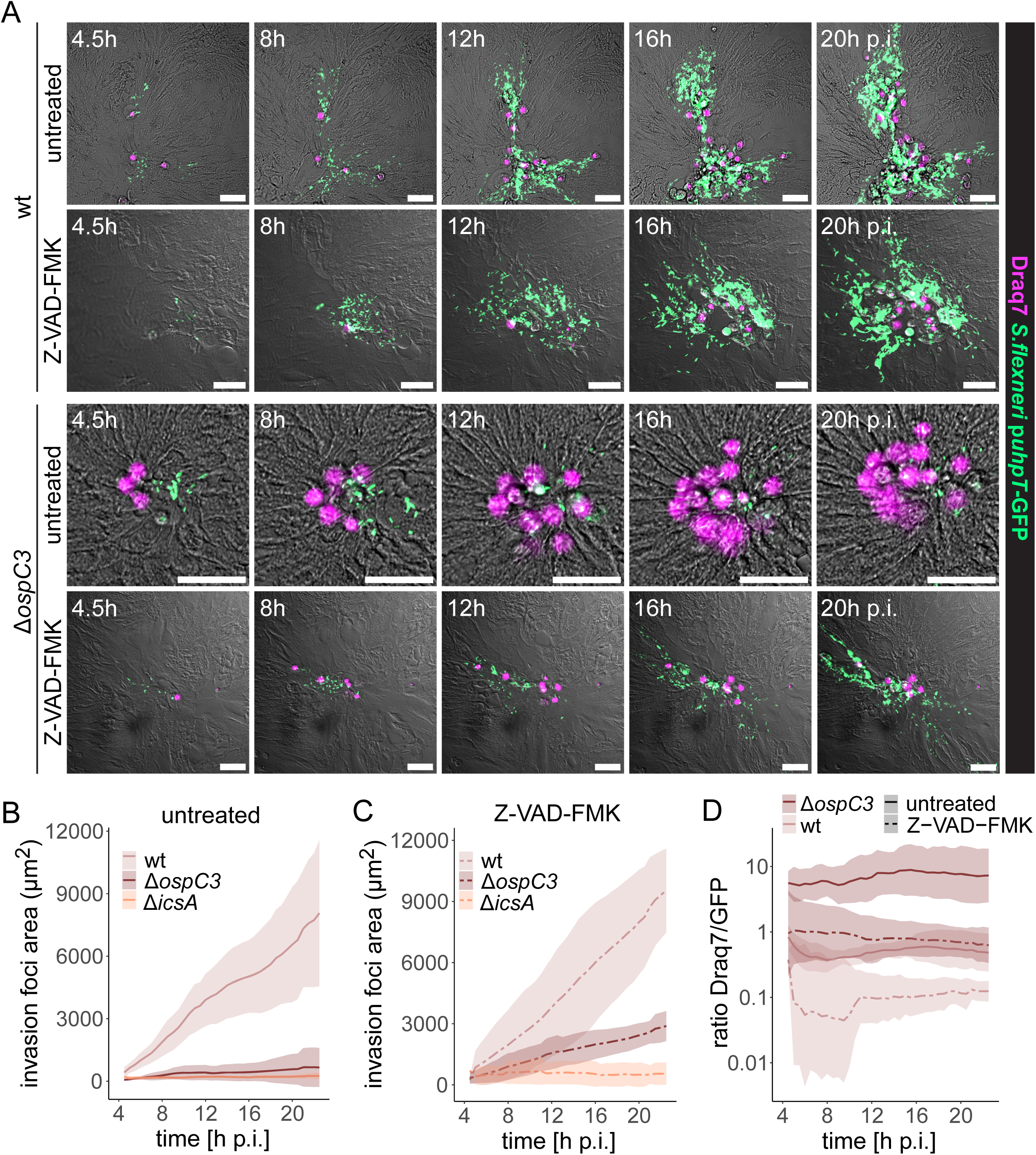
OspC3 inhibits caspase-dependent cell death to allow for expansion of the *Shigella* population within the colonoid epithelium. (A-D) Colonoid-derived monolayers were infected with the indicated *Shigella* p*uhpT*-GFP strains in the presence or absence of the broad-spectrum caspase inhibitor Z-VAD-FMK as in Figure 5 and individual invasion foci were followed by time-lapse microscopy. (A) Representative images of Z-VAD-FMK treatment restoring *Shigella* Δ*ospC3* intraepithelial expansion to wt levels. Scale bars: 50µm. Quantification of the GFP-positive area in (B) untreated and (C) Z-VAD-FMK treated monolayers over time. (D) The Draq7-to-GFP ratio for *Shigella* Δ*ospC3* was reduced to wt levels upon Z-VAD-FMK treatment. Data is plotted as mean + SD of 4-5 replicates per strain and treatment (2 replicates for Δ*icsA*, Z-VAD-FMK), with one replicate corresponding to the mean of 2-8 foci in an individually infected well.

**Supplementary Figure S6.**
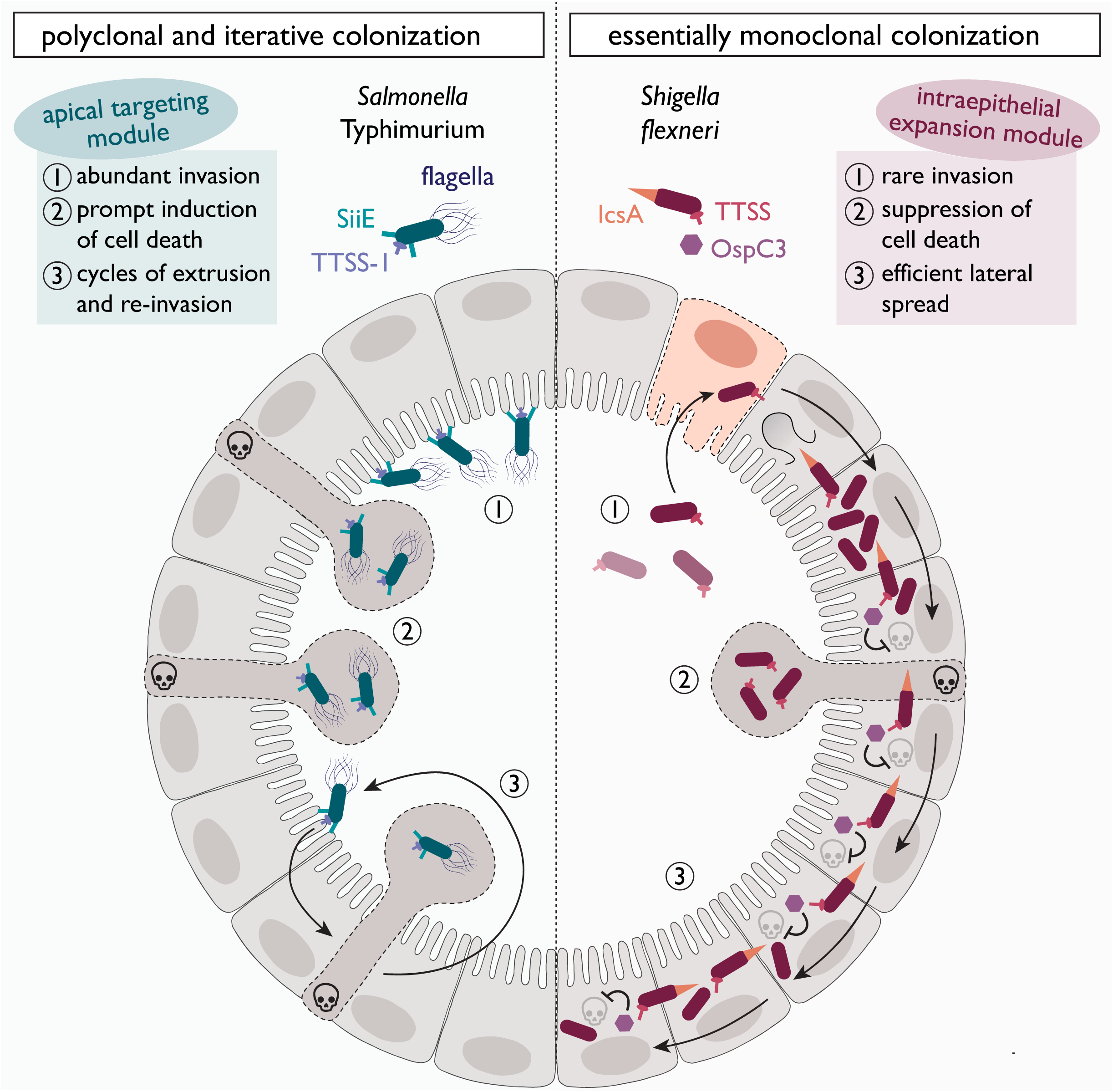
A conceptual model highlighting the basis for divergent *Salmonella* and *Shigella* colonization strategies of human intestinal epithelia. In the *Salmonella* polyclonal and iterative colonization strategy (left side), flagellar motility and the SPI-4-encoded adhesin system cooperate with the T3SS-1 to form an efficient apical targeting module. This results in efficient scanning of the epithelial surface for suitable binding sites and abundant invasion foci. However, these invasion events are typically abrogated by the prompt induction of IEC death, resulting in fast and iterative cycles of IEC invasion and extrusion. The *Shigella* essentially monoclonal colonization strategy (right side), on the other hand, is dependent on an intraepithelial expansion module comprising the inflammasome-inhibitory T3SS effector OspC3 and the actin nucleator IcsA. The concerted action of these effectors allows *Shigella* to laterally outrun the IEC death response and compensate for exceptionally rare invasion events by efficient intraepithelial expansion.

### SUPPLEMENTARY TABLES

**Table S1.** Strains and plasmids used in this study.

| Strain | Genotype | Reference |
| --- | --- | --- |
| *S.*Tm wt | SL1344, wt (SB300, Sm<sup>R</sup>) | (1) |
| *S.*Tm $\Delta invG$ | SL1344, $\Delta invG$ (SB161, Sm<sup>R</sup>) | (2) |
| *S.*Tm $\Delta SPI-4$ | SL1344, $\Delta SPI-4$ (Sm<sup>R</sup>, Kan<sup>R</sup>) | (3) |
| *S.*Tm $\Delta fljBfliC$ | SL1344, $\Delta fljBfliC$ (Sm<sup>R</sup>, Kan<sup>R</sup>) | (4) |
| *S.*Tm $\Delta fljBfliC \Delta invG$ | SL1344, $\Delta fljBfliC \Delta invG$ (Sm<sup>R</sup>, Kan<sup>R</sup>) | This study |
| *S.*Tm $\Delta fljBfliC \Delta SPI-4$ | SL1344, $\Delta fljBfliC \Delta SPI-4$ (Sm<sup>R</sup>, Kan<sup>R</sup>) | This study |
| *S.*Tm $\Delta motA \Delta SPI-4$ | SL1344, $\Delta motA \Delta SPI-4$ (Sm<sup>R</sup>, Kan<sup>R</sup>) | This study |
| *S. flexneri* wt | M90T, wt | (5) |
| *S. flexneri* $\Delta mxiD$ | M90T, $\Delta mxiD$ (Kan<sup>R</sup>) | (6) |
| *S. flexneri* $\Delta ospC3$ | M90T, $\Delta ospC3$ (Kan<sup>R</sup>) | This study |
| *S. flexneri* $\Delta icsA$ | M90T, $\Delta icsA$ (Kan<sup>R</sup>) | This study |
| Plasmid | Description | Reference |
| pFPV-mCherry | *rpsM*-mCherry (constitutive) | (7) |
| pM975 | *pssaG*-GFP (SPI2-dependent, vacuolar) | (8, 9) |
| pZ1400 | *puhpT*-GFP (cytosolic) | (10) |

**Table S2.**
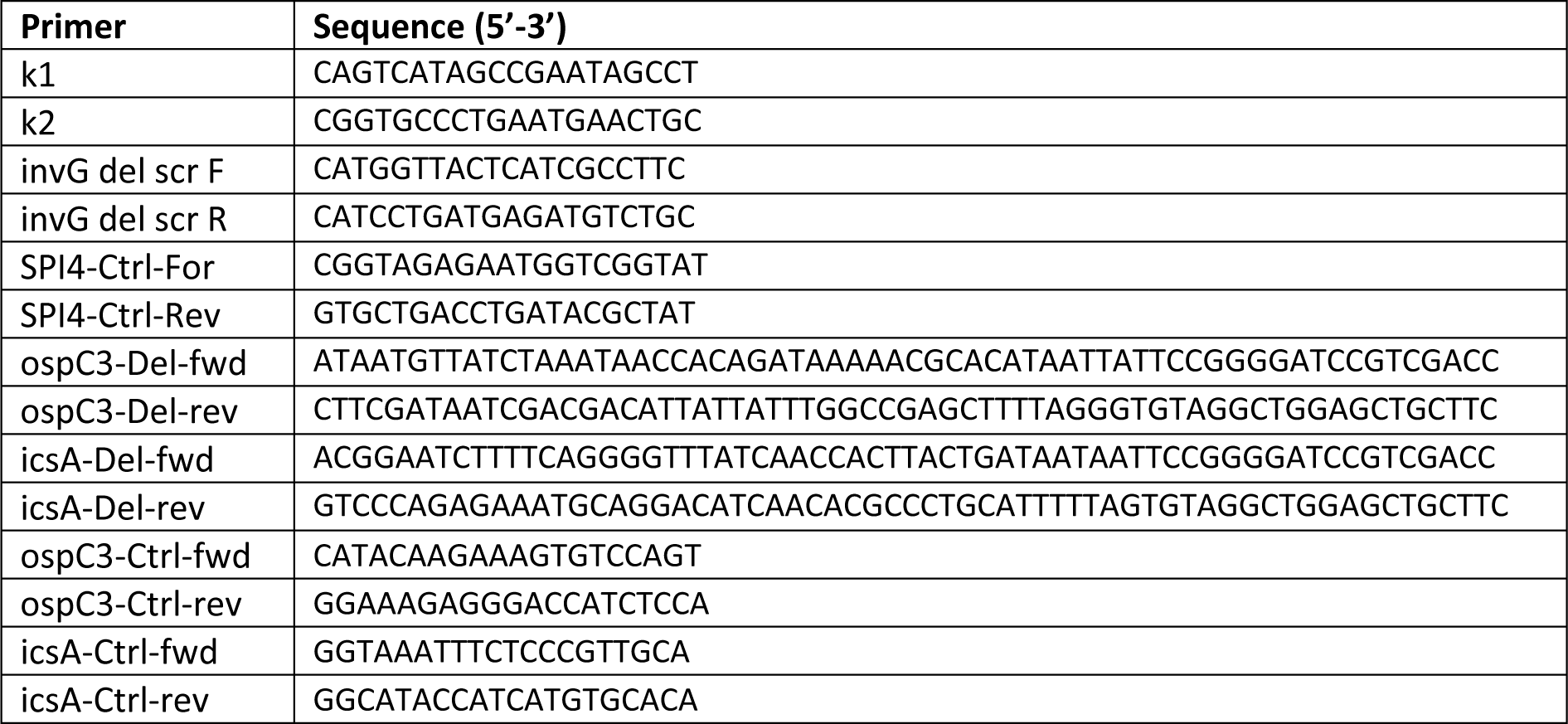
Primers used in this study.

## Notes

### Competing Interest Statement

The authors have declared no competing interest.

